# Co-targeting the PI3K-Akt pathway improves response to MEK inhibition in low-grade serous ovarian cancer cell lines

**DOI:** 10.1101/2025.09.05.674550

**Authors:** Rebekah M. Peplinski, Jesse D. Riordan, Jacob L. Schillo, Elizabeth C. Hannan, Silvana Pedra Nobre, Yasmin A. Lyons, Keely K. Ulmer, Michael J. Goodheart, Adam J. Dupuy

## Abstract

Low grade serous ovarian cancer (LGSOC) accounts for ∼5% of all ovarian cancers, with >1,000 new cases diagnosed each year in the United States. Currently, there is no established standard-of-care treatment for this subtype, and chemotherapy response rates are only ∼4%. One defining characteristic of LGSOC is a high prevalence of mutations within the MAPK pathway, which has motivated studies using MEK inhibitors to treat this disease. A recent phase II/III clinical trial using trametinib to treat patients with recurrent LGSOC improved the response rate to 26%; however, all patients eventually progressed due to the emergence of therapeutic resistance. Ongoing clinical trials are investigating combined inhibition of MEK and FAK with avutometinib plus defactinib for LGSOC patients, with preliminary results demonstrating response rates near 50%. While these results are highly encouraging, a significant subset of patients still fails to respond to existing targeted treatments. Additionally, development of therapeutic resistance in initial responders remains a major obstacle. Here, we sought to characterize molecular responses of LGSOC to MEK inhibition, both acutely and during the development of spontaneous drug resistance, with a goal of identifying adaptive mechanisms LGSOC cells utilize to withstand MAPK inhibition. Our research identified significant upregulation of the PI3K-Akt pathway in response to MEK inhibition. While targeted inhibition of AKT had minimal impact on its own, combination with MEK inhibitors produced strong synergistic suppression of proliferation in LGSOC cells. This combination strategy could potentially be used to prevent or reverse the emergence of MEK inhibitor resistance in LGSOC patients.

## Introduction

Low-grade serous ovarian cancer (LGSOC) has recently been characterized as a unique subtype of ovarian cancer with distinct molecular characteristics, clinical treatments, and therapeutic responses [1]. These major characteristics include a low mitotic rate, wild-type p53 status, and frequent mutations within the mitogen-activated protein kinase (MAPK) pathway [2]. Its counterpart, high-grade serous ovarian cancer (HGSOC) has a high mitotic rate, mutant *TP53*, and frequent mutations in BRCA1/2 DNA repair associated (*BRCA1/2*), checkpoint kinase 2 (*CHEK2*), and RAD51 recombinase paralog C (*RAD51C*) (DNA repair mutations) [2], making it physiologically and molecularly distinct from LGSOC.

LGSOC accounts for ∼5% of epithelial ovarian cancers, and ∼12,000 women in the United States are currently living with this disease. It disproportionally affects younger women, with a median age of diagnosis around 45 years. As compared to HGSOC, LGSOC is associated with slower progression and prolonged survival but a poor quality of life. Given its slow-progressing nature and earlier onset, women can live with the disease for longer periods of time, increasing morbidity. LGSOC has <5% overall response rate to current chemotherapy, and due to a historical lack of effective treatments, affected women experience years of symptomatic disease with few (if any) disease-free intervals, leading to an overall poor quality of life [3-8].

The high prevalence of MAPK pathway genetic alterations in LGSOC (>60% overall; ∼30% Kirsten Rat Sarcoma virus (*KRAS*), ∼10% Neuroblastoma Rat Sarcoma virus (*NRAS*), ∼10% B-Raf proto-oncogene (*BRAF)* ∼10% other MAPK pathway genes) [2, 9, 10] has led to recent MAPK-targeted therapeutic clinical trials (NCT02101788, NCT04625270). Mitogen-activated protein kinase kinase (*MEK1/2*) inhibition, particularly with trametinib, has been shown to be effective in patients with recurrent LGSOC, increasing the response rate to 26%, as compared to 6% on physicians’ choice (progression-free survival of 13 and 7.2 months, respectively) [11]. This improved response rate suggests targeting MEK as a potential first-line therapy to treat LGSOC. However, patients that respond initially eventually progress due to acquired therapeutic resistance, with no remaining effective treatment options. Here, we present a series of experiments characterizing molecular responses of LGSOC to MEK inhibition in a panel of established and patient-derived cell lines. We identified multiple candidate mechanisms that may be targeted in combination to improve therapeutic efficacy and outcomes for LGSOC patients. Among these candidates, we validated and characterized the synergistic impact of co-targeting the PI3K/Akt signaling pathway to enhance antiproliferative effects of MEK inhibition in LGSOC cells.

## Results

### LGSOC cell lines are sensitive to trametinib and spontaneous resistance confers reduced MAPK dependence

A panel of cell lines, genetically characterized as LGSOC [12, 13], was treated with a range of trametinib doses to establish a dose-response curve and confirm sensitivity (Fig 1A). The patient-derived cell line (VOA-7681) was the most sensitive of the cell lines, followed by the established cell lines OV-56, ES-2, and OVCAR-8. This varying sensitivity across the panel allows us to study the effects of trametinib at different degrees of responsiveness—much like patient responses. Western blot analysis showed a strong decrease in phosphorylated extracellular signal-regulated kinase (p-ERK) levels—the protein kinase downstream of MEK in the MAPK pathway—with trametinib treatment for all four cell lines, confirming inhibition of the MAPK pathway (Fig 1B). After 2-3 weeks of trametinib exposure at an IC_95_ dose, these cell lines develop spontaneous resistance, with ES-2 having the most robust response. After an initial drop in proliferation, development of spontaneous resistance returned ES-2 cells to normal growth rate under constant trametinib treatment, with an increase in the IC_50_ from 4.5 nM (parental) to 135.5 nM (trametinib-resistant) (Fig 1C). Despite regained proliferative capacity, p-ERK levels do not return to baseline in resistant cells (Fig 1B), indicating continued inhibition of the MAPK pathway. However, compared to cells treated for 24 hours, a low level of p-ERK does return in cells treated with a constant dose of trametinib for three weeks (Fig 1B), indicating a low level of MAPK pathway reactivation.

**Figure 1.**
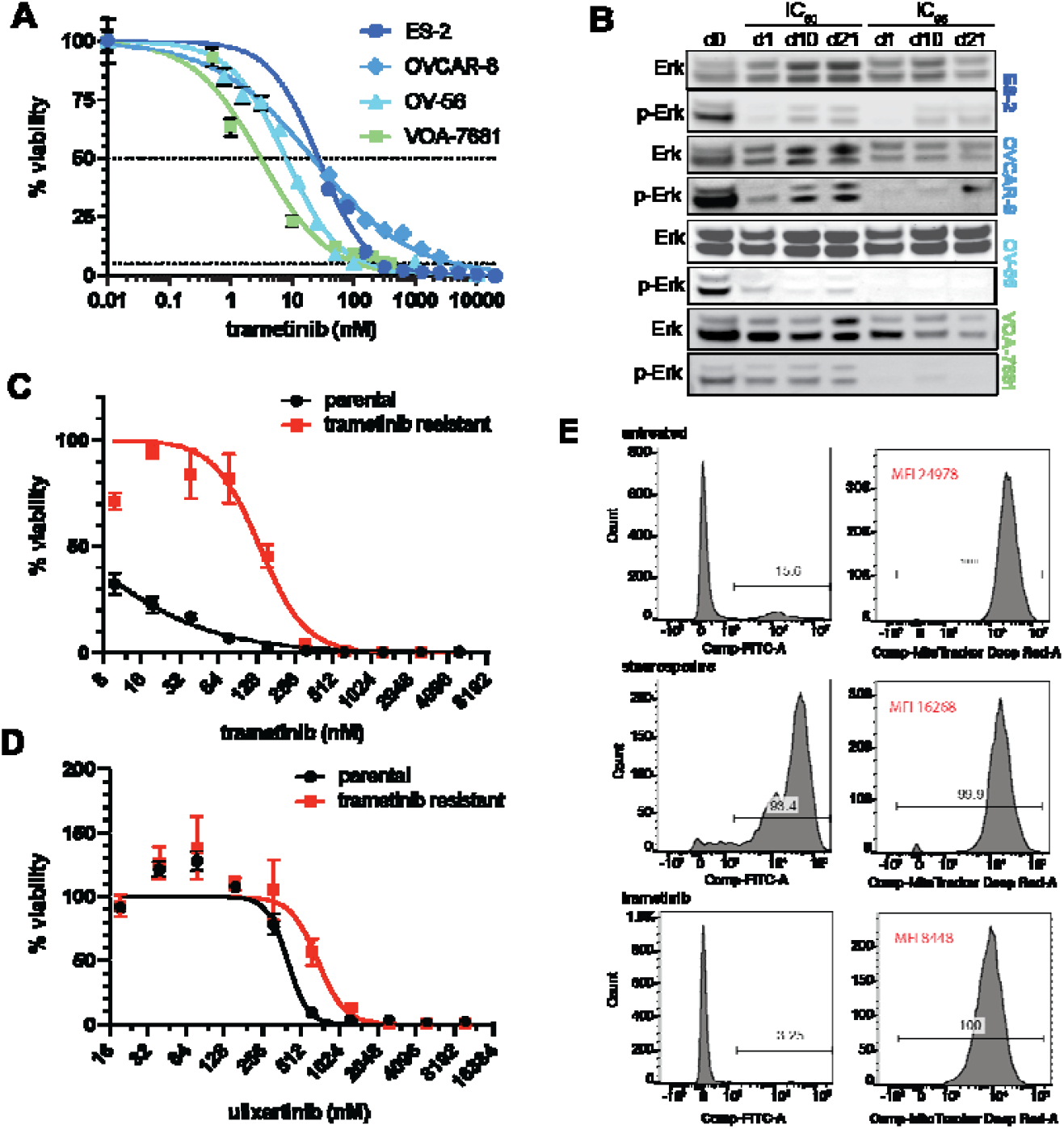
LGSOC cell lines are sensitive to trametinib in vitro. **(A)** Dose-response curves showing sensitivity to trametinib as percent viability double-normalized to day 0 and untreated cells. Dotted lines show IC_50_ (top) and IC_95_ (bottom). **(B)** Western blot analysis showing a decrease in p-ERK over a three week time-course. **(C)** Dose response curves for parental and trametinib resistant ES-2 cells treated with trametinib confirming resistance. **(D)** Dose response curves for parental and trametinib resistant cells treated with ulixertinib to determine the role of reactivation of pERK in resistance. **(E)** Flow for annexin V (left; FITC— apoptosis) and Cytopainter (right; DeepRed—cell proliferation) showing minimal cytotoxicity and inhibition of proliferation. Untreated cells were stained with CytoPainter 30 min prior to flow. Cells were treated with staurosporin (1µM) for 24 hours to induce apoptosis or trametinib (20nM) for 7 days.

To determine the impact of remaining MAPK pathway activity on proliferation, we treated trametinib-resistant cells with the ERK inhibitor ulixertinib. Compared to the parental cells, trametinib-resistant cells had a significantly increased ulixertinib IC_50_ (400 nM vs 688 nM for ES-2 cells; Fig 1D), indicating that the resistant cells may be less reliant on the reactivated MAPK pathway, though not completely ERK-independent. Additionally, annexin V staining following trametinib treatment showed low levels of apoptotic cells, indicating an anti-proliferative rather than cytotoxic effect (Fig 1E). Taken together, our results suggest that trametinib induces strong inhibition of downstream MAPK pathway signaling initially in LGSOC cells, which is associated with impaired proliferation but not induction of apoptosis. As resistance to trametinib emerges, cells reactivate ERK signaling at low levels and demonstrate reduced but not absent dependency on this signaling pathway.

### CRISPR kinome knockout screen identified several protein kinases functionally relevant to trametinib treatmen

Abnormal kinase activity is commonly implicated in cancer, and protein kinases are the second most commonly targeted class of proteins after G-protein coupled receptors [14]. A focused CRISPR screen was used to individually induce deleterious mutations in 763 human kinase genes [15] to determine which targets produce a synthetic lethality phenotype in the context of low-level trametinib treatment. Synthetic lethality occurs when targeting two genes or pathways simultaneously leads to cell death, while the loss of either gene or pathway alone is tolerated. We chose to conduct kinome-targeted CRISPR screens in ES-2 and OVCAR-8 cell lines due to their relatively higher trametinib IC_50_ values (Fig 1B, Fig S1). The effect of low dose trametinib treatment on these cell lines should be minimal, thus providing increased statistical power to observe synthetic lethality with individual kinase deletions (*i*.*e*., decreased abundance of specific guides in the population due to dropout of cells harboring them). The remaining trametinib-sensitive lines (VOA-7681 and OV-56) were used for validation studies alongside ES-2 and OVCAR-8.

ES-2 and OVCAR-8 cells were transduced with the lentiviral Brunello kinome sub-libraries [15] containing Cas9 and sgRNAs at MOI 0.5 to ensure that most cells receiving sgRNAs got only one guide targeting a single kinase. After selection for stable integration of sgRNA constructs, cells were treated with either vehicle for four days or trametinib for ten days using a dose of trametinib that controlled growth so that cells were collected at the same confluency. This strategy was used to ensure that each population underwent an equal number of cell doublings prior to collection (Fig S2). The kinome sgRNA library was then amplified from each cell population using simple PCR techniques and sequenced to identify the relative abundance of each guide in each population [15]. We then utilized the Model-based Analysis of Genome-wide CRISPR/Cas9 Knockout (MAGeCK) method for prioritizing candidate genes exhibiting positive or negative selection for knockout in the vehicle-treated and drug-treated populations based on significant enrichment or depletion, respectively, of guides targeting them [16]. Significant depletion of multiple independent guides targeting specific kinases in the vehicle- and trametinib-treated populations after outgrowth identifies “essential” kinases that are necessary for survival regardless of treatment status, such as those regulating general cell cycle progression. Depletion or dropout of guides only in trametinib-treated but not vehicle-treated populations identifies kinases important for survival under trametinib treatment, while specific enrichment of guides in these populations identifies kinases that could promote trametinib resistance. Enriched sgRNAs specific to trametinib-treated populations included guides targeting genes involved in BMPR and TGFβ signaling, suggesting inactivation of these pathways may promote resistance. (Table S1). Kinases of highest interest were those whose guides dropped out only in the presence of trametinib, indicating a synthetically lethal combination (Fig 2A). Among these synthetically lethal hits were kinases within the MAPK and phosphatidylinositol 3-kinase (PI3K)/protein kinase B (AKT) pathways (Table S1). Western blot analysis confirmed trametinib-induced upregulation of phosphorylated PI3K (p-PI3K) and phosphorylated AKT (p-AKT) in independent LGSOC cell populations, indicating that the PI3K-Akt pathway is activated in response to trametinib treatment (Fig 2B-C).

**Figure 2.**
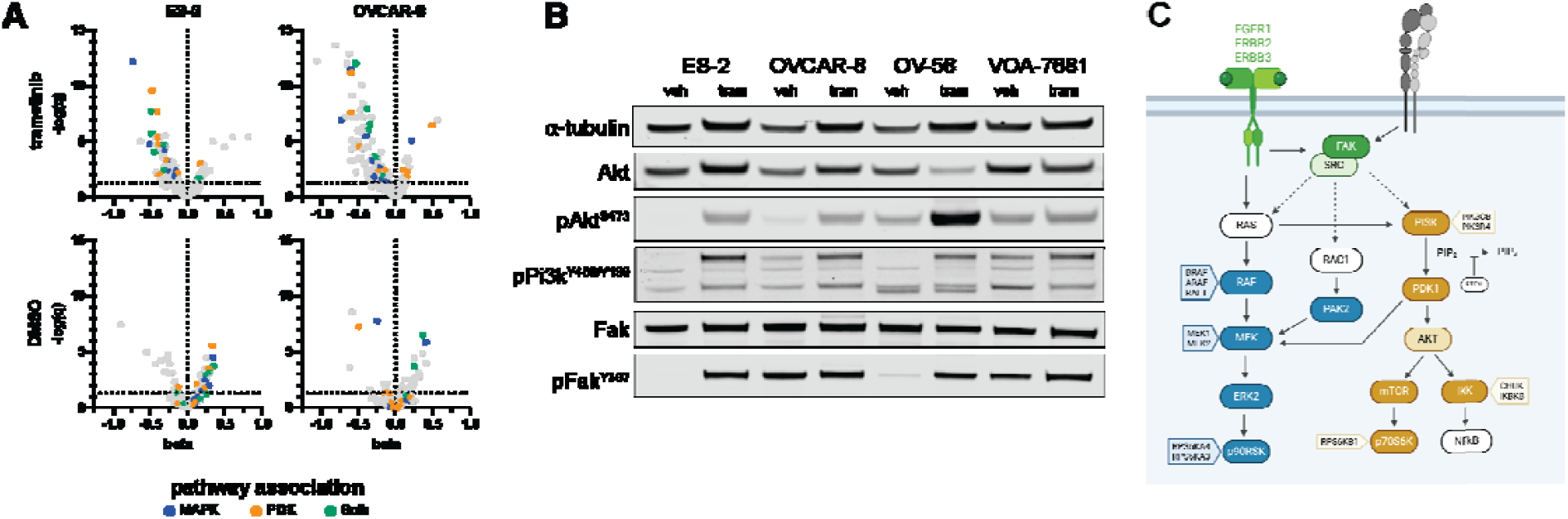
Identification of synthetic lethal interactions in LGSOC cell lines *in vitro* using a CRISPR kinome library. **(A)** Volcano plots showing log-fold-change (beta) and significance (-log(q)) of sgRNAs from ES-2 and OVCAR-8 screens in both trametinib-treated and vehicle-treated populations. Highlighted sgRNAs are associated with the MAPK pathway (blue), PI3K-Akt pathway (orange), or both pathways (green). **(B)** Western blots confirming increase in PI3K-Akt pathway signaling under trametinib treatment. **(C)** Working model for signaling through the MAPK and PI3K-Akt pathways under trametinib treatment.

### Trametinib paired with AKT inhibitor capivasertib synergistically inhibits LGSOC cell proliferation

Our CRISPR screen identified several kinases as candidates whose inhibition may enhance the antiproliferative effects of trametinib in LGSOC cells, many of which have inhibitors that are either currently being used in clinical trials or have been FDA-approved for other cancer types. To identify synergistic or additive drug effects, we focused on MAPK and PI3K-Akt pathway inhibitors, prioritized based on our kinome screen results and drug availability for clinical use. The LGSOC cell line panel was treated with various combinations of trametinib (MEKi), avutometinib (RAF/MEKi), VS-4718 (FAKi; surrogate for defactinib used in preclinical models), and capivasertib (AKTi) to assess effects on cell proliferation and apoptosis. Avutometinib and VS-4718 were chosen based on a recent phase II clinical trial showing a 46% response rate when avutometinib and defactinib were used in combination to treat recurrent LGSOC, which was published as this work was being completed [17]. Notably, activated FAK can signal to both MAPK and PI3K-Akt pathways. While we did identify protein tyrosine kinase 2 (*FAK*) as a candidate synthetic lethal kinase in our CRISPR screen, it was not one of the strongest hits (q= 4.59E-06 (ES-2) and 0.0036368 (OVCAR-8)) (Table S1). In comparison, 3-Phosphoinositide Dependent Protein Kinase 1 (*PDPK1*) – a kinase in the PI3K-Akt pathway— was one of the top hits (q= 2.44E-10 (ES-2) and 6.69E-12 (OVCAR-8) (Table S1). Additionally, phosphorylated FAK (Tyr397) (p-FAK)—the site of phosphorylation via PI3K—was increased upon trametinib treatment in 3 of 4 LGSOC cell lines (Fig 2B), indicating that FAK signaling may be mechanistically involved in trametinib response. This result is consistent with a model in which MEK inhibition leads to activation of the PI3K-Akt pathway downstream of FAK (Fig 2C).

Taking the recent success of avutometinib and defactinib combination therapy in LGSOC into account [17, 18], we used this condition as a basis of comparison for LGSOC cell responses to treatment with combined MAPK and PI3K/AKT pathway inhibitors. To investigate synergistic relationships, we treated our panel of cell lines with escalating doses of specific paired drug combinations—avutometinib plus VS-4718, trametinib plus capivasertib, or avutometinib plus capivasertib. This experimental strategy creates a grid-like matrix testing the effectiveness of each drug within a pair at multiple concentrations of the other drug, using cell viability as the readout. Synergy scores were calculated using SynergyFinder+ [19]. The synergy scores determined for each cell line indicated a strong synergistic effect (scores > 10) between trametinib and capivasertib compared to either of the avutometinib combinations, which would instead be considered additive (scores from -10 to 10) (Fig 3A-D, Table S2). Annexin V staining following treatments in the VOA-7681 cell line indicated that both combination treatments (trametinib/capivasertib or avutometinib/VS-4718) are anti-proliferative, rather than cytotoxic (Fig 3E). Western blot confirmed that the drugs used were hitting their intended targets (Fig 3F), with a decrease in p-ERK in response to MEK inhibition, decrease in p-FAK in response to VS-4718, and an increase in p-AKT in response to capivasertib—indicative of an accumulation of sequestered, active AKT.

**Figure 3.**
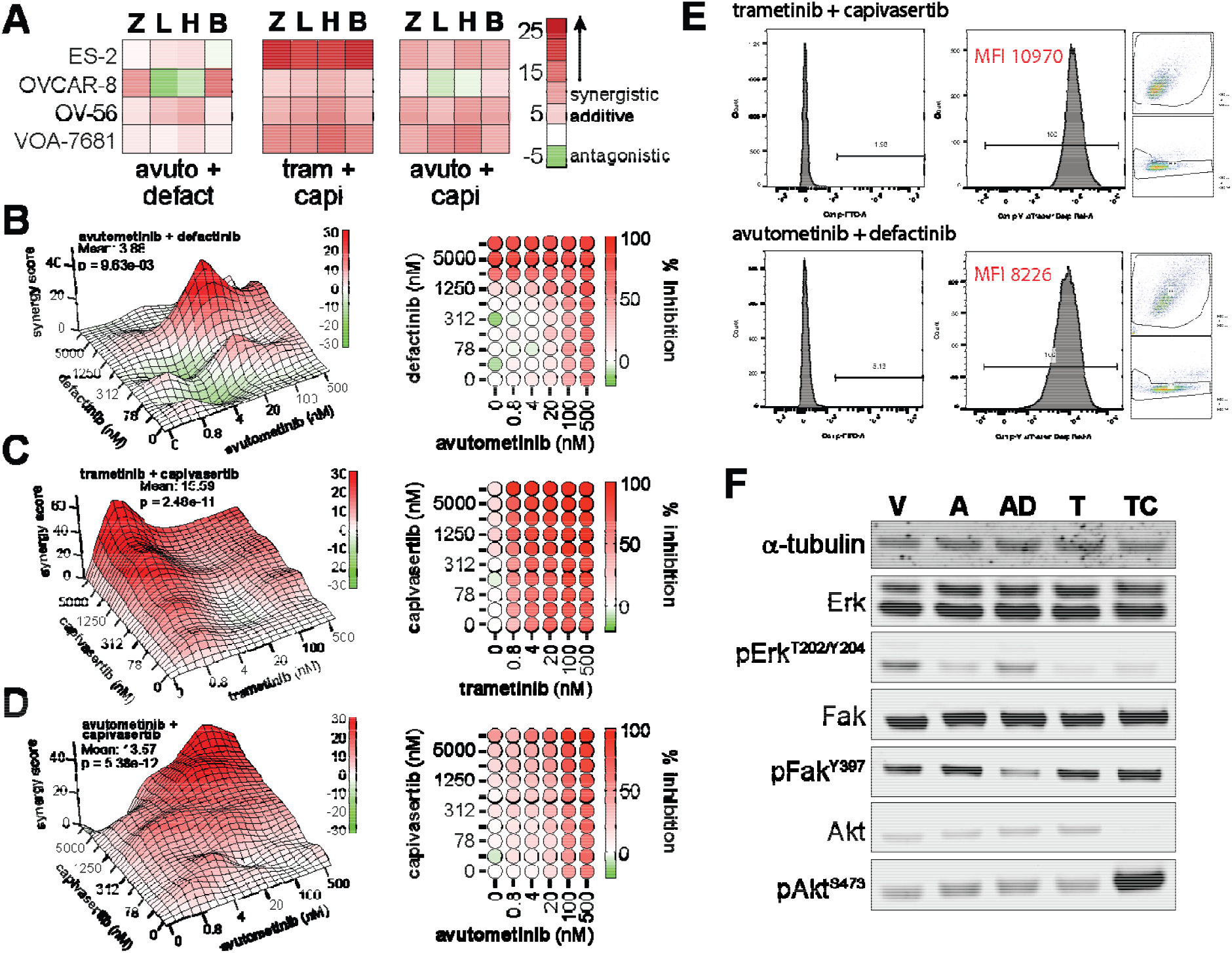
Identification of effective combination therapies to treat LGSOC *in vitro*. **(A)** Synergy scores for each cell line (Z=ZIP, L= Loewe, H=HSA, B=Bliss). **(B-D)** 3D plots showing synergy scores (left) and heat maps showing percent inhibition for: avutometinib and defactinib (B), trametinib and capivasertib (C), and avutometinib and capivasertib (D). **(E)** Flow for annexin V (Left; FITC—apoptosis) and Cytopainter (Right; DeepRed—cell proliferation) showing minimal cytotoxicity and an inhibition of proliferation. Cells were treated with trametinib (20nM) and capivasertib (3.2µM) or avutometinib (750nM) and VS-4718 (535nM) for 7 days. Untreated cells and staurosporin-treated cells are shown in Figure 1D. **(F)** Western blot analysis confirm on-target activity of drugs. Therapeutic doses used are the same as the prior panel (V, vehicle; A, avutometinib; AD, avutometinib/defactinib; T, trametinib; TC, trametinib/capivasertib). Upregulation of pAkt under capivasertib treatment is indicative of inhibition via sequestration.

## Discussion

There is a critical need to understand the biology of oncogenic MAPK signaling in LGSOC and its key mechanistic mediators. Such understanding will drive the development and advancement of molecular targeted therapies to improve patient responses and outcomes, as has been the case for several other cancer types. For example, after mutations in *anaplastic lymphoma kinase* (*ALK)* were identified as primary oncogenic drivers in some lung cancers, lorlatinib (an ALK inhibitor) improved long-term response rate from 15% to 74%, extending 5-year progression-free survival from 8% to 60% [20]. Treatment of non-small cell lung cancers and cutaneous melanomas harboring *BRAF*^*V600E*^ mutations with targeted BRAF and MEK inhibitors encorafenib plus binimetinib controlled disease in 75% and 80% of patients, respectively [21, 22]. Based on these and other successful examples, improved understanding of the biology and molecular mechanisms underlying oncogenic signaling pathways and their responses to therapeutic inhibition could lead to improved efficacy and duration of therapeutic response in LGSOC patients.

Recognition of LGSOC as a distinct entity from HGSOC with frequent MAPK pathway-activating mutations raised hopes that targeting this pathway would significantly improve patient outcomes. While some success has been reported and other clinical trials are ongoing, improvements to date have been modest in both magnitude and duration. The first trial with a MEK inhibitor, selumetinib, reported a 15% overall response rate and did not proceed past Phase II [23]. A subsequent trial with a different MEK inhibitor, binimetinib, had a similar response rate of 16% and failed to extend progression free survival, as compared to standard chemotherapy [24]. More recent MEK inhibitor trials have demonstrated improved success in LGSOC, with overall response rates of 26% and 46% for trametinib and avutometinib, respectively [11, 17]. While encouraging, a majority of patients remains unresponsive to these treatments. Furthermore, patients that do respond initially often develop therapeutic resistance and progressive disease with no remaining treatment options. Taken together, these clinical trial results indicate that although activated MAPK signaling is a key oncogenic driver in LGSOC, targeted inhibition of MEK alone does not adequately eliminate disease or ultimately prevent progression. More work must be done to understand the key mechanistic regulators of oncogenic MAPK signaling in LGSOC, as well as genes and pathways that cooperate with and/or compensate for MAPK signaling when it is targeted therapeutically. Unfortunately, molecular studies of LGSOC are highly limited and rarely extensive, in part due to its relatively recent unique classification from HGSOC and comparative rarity. Some studies have begun to investigate combinatorial treatment strategies and therapeutic resistance mechanisms. Clinically, inhibitors for CDK4/6, PIK3CA, MEK, and FAK are being investigated in various combinations with or without hormonal therapy [25]. A recent preclinical study implicated NOTCH signaling in acquired MEK inhibitor resistance and suggested that targeting MEK in combination with a pan-RAF inhibitor could be an effective treatment for LGSOC [26]. Here, we used similar preclinical approaches to uncover the biology behind oncogenic MAPK signaling in LGSOC and identified PI3K-Akt signaling pathway activity as a synthetic lethal vulnerability in the context of MEK inhibition with trametinib. PI3K-Akt signaling was strongly upregulated in response to MEK inhibition (Fig. 2B), and targeted inhibition of AKT synergized with MEK inhibition to impair LGSOC cell proliferation (Fig. 3).

We selected a panel of LGSOC cell lines based on their molecular profiles [12, 13] and confirmed trametinib sensitivity, consistent with an activated MAPK-signaling pathway phenotype (Fig 1A). While p-ERK levels decreased dramatically in trametinib treated cells—confirming inhibition of the MAPK pathway— spontaneous resistance emerged within 21 days of treatment, with p-ERK levels increasing slightly but not returning to baseline (Fig 1B). Dose-response curves confirmed an extreme increase in trametinib IC_50_ in emergent populations compared to parental cells in ES-2 (Fig 1C). In other cancer types, such as cutaneous melanoma, p-ERK levels typically return to baseline after initially decreasing during the spontaneous development of MAPK pathway inhibitor resistance, indicating full reactivation of MAPK signaling [27-32]. Trametinib-resistant LGSOC cells didn’t fully restore p-ERK levels, and their IC_50_ for the ERK inhibitor ulixertinib was significantly increased compared to parental cells (Fig 1D), suggesting resistant cells are less reliant on MAPK signaling and may instead use alternative pathways to maintain proliferation and survival. Additionally, annexin V staining showed that trametinib does not strongly induce apoptosis, indicating that the effect of trametinib is antiproliferative, rather than cytotoxic (Fig1E). Taken together, these results demonstrate that LGSOC cells can develop spontaneous resistance to MEK inhibitor-induced cytostasis in a manner that, while still ultimately dependent on some level of p-ERK activity, likely involves MAPK-pathway-independent mechanisms.

To identify functionally relevant genes and pathways involved in the response of LGSOC cells to MEK inhibition, we utilized a targeted CRISPR screening approach paired with trametinib treatment. This approach allowed us to identify kinases whose loss prevented outgrowth only in treated cells, indicating a synthetic lethal interaction with trametinib. These kinases represent candidates that could potentially be targeted for inhibition in combination with trametinib to increase its effectiveness and/or prevent therapeutic resistance. Given that many drugs that are FDA approved or in clinical trials target kinases, this approach has strong potential to identify effective combinations of drugs that are already available in the clinic, enhancing the potential for rapid translational impact. We performed CRISPR kinome screens in the two most resistant cell lines of our LGSOC panel (ES-2 and OVCAR-8) to identify candidate kinases (Table S1), finding good reproducibility across biological replicates, particularly among strongly enriched or depleted guide RNAs (Fig S1). Many kinases identified from this screen with significantly depleted guide RNAs are members of the MAPK and PI3K-Akt pathways, including FAK, RAF, PI3K, PDPK1, and AKT substrates (IKKα and IKKβ) (Fig 2A-B). Targeting these kinases, most of which have inhibitors currently used in the clinic, could improve patient response to MEK inhibition. Given the reemergence of low levels of p-ERK in cells treated with trametinib long-term, there could be some reactivation of the MAPK pathway that contributes to regained proliferative status. This is a common resistance mechanism seen in other cancers with frequent MAPK mutations like cutaneous melanoma [27-32] and could explain why kinases within the MAPK pathway were identified as candidates by our screen. Although AKT was not identified as a strong candidate in our screen directly– likely due to redundancy amongst the homologous isoforms—we did identify several kinases within the PI3K-Akt pathway, including AKT substrates IKKα and IKKβ, thereby functionally implicating PI3K-Akt signaling in trametinib resistance (Fig 2, Table S1). Additionally, PDPK1 was a strong hit identified from our screen which has multiple targets including AKT, protein kinase C (PKC), ribosomal protein S6 kinase (RPS6K), and serum/glucocorticoid regulated kinase (SGK). It is possible that one or more of these other PDPK1 targets plays a role in MEK inhibitor resistance by activating alternative pathways the cells can use to overcome MAPK pathway inhibition.

While this work was being completed, avutometinib and defactinib combination treatment was granted FDA orphan drug designation for treating recurrent LGSOC [17]. Although loss of FAK, the primary target of defactinib, did exhibit synthetic lethality with trametinib treatment in our CRISPR screen, it wasn’t as strong of a hit as other kinases within the MAPK or PI3K-Akt pathways (Table S1). Additionally, western blot showed that baseline p-FAK levels varied across LGSOC cell lines and were increased in some but not all in response to trametinib treatment (Fig 2B). Variable utilization of FAK signaling in trametinib-treated patients could explain in part why the response rate for avutometinib and defactinib was suboptimal (46%) [17]. p-PI3K and p-AKT were more consistently increased by trametinib treatment across the cell panel (Fig 2B). This observation indicates that LGSOC cells upregulate PI3K signaling in response to MEK inhibition.

Based on drug synergy assays, the combination of trametinib and capivasertib outperformed the avutometinib and defactinib combination across all four tested LGSOC cell lines (Fig 3, Table S2). Importantly, avutometinib plus capivasertib also performed better than avutometinib plus defactinib. This result could indicate that FAK is just one way to ultimately activate the PI3K-Akt pathway and that inhibiting this pathway at a more critical downstream level (i.e., directly targeting Akt) results in more effective growth inhibition. Another potential explanation is that the MAPK and PI3K-Akt pathways are acting in parallel with separate functions within the context of LGSOC.

Additionally, based on the results from this study showing an increase in PI3K-Akt signaling in response to trametinib treatment, phosphatidylinositol 3-kinase (PTEN) loss or phosphatidylinositol-4,5-bisphosphate 3-kinase catalytic subunit alpha (PIK3CA) mutations, seen in a small subset of ovarian cancer patients [9, 10], could be a predictor of poor MEK inhibitor response (Fig. 2C). PTEN is a negative regulator of the PI3K-Akt pathway, and PIK3CA encodes the catalytic subunit of PI3K. Loss of PTEN or activating mutations in PIK3CA would lead to constitutive activation of the PI3K-Akt pathway, which could lead cells to rely heavily on the PI3K-Akt pathway to maintain cell growth and proliferation rather than the MAPK pathway, thereby decreasing MEK inhibitor effectiveness. This mutational status should be considered when deciding treatment strategies for patients.

There is a critical need in the ovarian cancer field to understand the biology behind therapeutic responses, their relative effectiveness, and the emergence of resistant tumors. Given that LGSOC is a relatively rare cancer and targeted therapy in LGSOC is a recent trend, there is currently a lack of matched pre- and post-treatment tumor samples to characterize the biology of this cancer directly in patient samples. Such matched biopsies would be an invaluable resource to identify potential resistance mechanisms. In their absence, we have utilized the available models of LGSOC [12, 13, 33] to identify candidate drivers of therapeutic resistance. As patient samples become available, targeted analyses can be conducted to determine whether the same mechanisms promote resistance in the clinic.

In summary, using a functional genomics approach, we have identified both MAPK and PI3K-Akt signaling pathways as major mediators of response to MEK inhibition in LGSOC. Loss of PI3K-Akt activity, either through CRISPR-induced mutation or targeted inhibition, strongly suppressed LGSOC cell proliferation and development of trametinib resistance. In drug synergy assays, Akt inhibitor capivasertib was more effective when paired with either avutometinib or trametinib than the clinically-approved combination of avutometinib plus defactinib. Based on these data, we suggest that combination treatment with trametinib plus capivasertib could provide improved patient outcomes and that further clinical study is warranted.

## Methods

### Cell lines

Cell lines were obtained from ATCC (ES-2; CRL-1978), NCI DCTD tumor repository (OVCAR-8), Millipore Sigma (OV-56; #96020759), and a generous donation from Dr. Mark Carey (VOA-7681). Cells were cultured at 37^°^C in 5% CO_2_ in the following media: McCoy’s 5A supplemented with 10% FBS and 1% Penicillin-Streptomycin (ES-2), RPMI supplemented with 10% FBS and 1% Penicillin-Streptomycin (OVCAR-8), DMEM/F12 supplemented with 5% FBS, 1% Penicillin-Streptomycin, 0.5µg/mL hydrocortisone (Targetmol T1614) and 10µg/mL bovine insulin (Sigma I0516) (OV-56), and MCDB105 and M199 (1:1) supplemented with 10% FBS and 1% Penicillin-Streptomycin (VOA-7681).

### Drugs

The drugs utilized throughout this work were obtained from Targetmol: trametinib (T2125), ulixertinib (T7005) capivasertib (T1920), avutometinib (T6971), and defactinib (T1996).

### Cell viability assay

Cells were plated at 750-1,000 cells/well in a 96-well plate. Drug-containing media was replenished every 3-4 days. To assess viability, resazurin sodium salt (Sigma–Aldrich, R7017) resuspended in Dulbecco’s phosphate-buffered saline was added to each well to a final concentration of 25µg/mL and then incubated at 37^°^C for 2 hours. Fluorescent signal (560nm excitation/590nm emission) was measured using the BioTek® Synergy™ HT microplate reader. Percent viability was calculated by double normalizing to day 0 read and vehicle treated control. Synergy scores were determined using the Synergy Finder+ web application [19].

### CRISPR kinome screen

Cells were plated at 1×10^6^ cells per plate in a 10cm plate. 24 hours later, cells were transduced with the Brunello kinome sub-libraries (Addgene #75314 and #75315) at an MOI of 0.5. Cells then underwent puromycin selection. Once selected, cells were plated in T175 flasks and treated for either 4 days (vehicle) or 10 days (trametinib). Both vehicle and trametinib treated cells were collected at equivalent confluency and treatment doses were selected based on 10-day dose response curve at ∼40% viability (Fig S2).

gDNA extractions were completed using Monarch® Genomic DNA Purification Kit (NEB, #T1310). sgRNA sequences were amplified by standard PCR techniques including 5µL 10X reaction buffer, 1.5µL 50mM MgCl_2_, 1µL 10mM dNTPs, 2µL 10µM P5 primer pool, 2µL 10µM P7 primer pool (see Table S3 for sequences), 1µg gDNA, 0.2µL Platinum Taq Polymerase, and 18.3μl H_2_O for a total volume of 50μl. Cycle conditions were as follows:

- Step 1: 94^°^C for 2 min.
- Step 2: 94^°^C for 30s, 55–65^°^C for 30s, and 72^°^C for 30s; repeat 15 cycles.
- Step 3: 72^°^C for 5 min; hold at 4^°^C.

Samples were then barcoded during the secondary PCR reaction including 5µL 10X reaction buffer, 1.5µL 50mM MgCl_2_, 1µL 10mM dNTPs, 1µL 10µM xGen UDI primers (IDT product #10008052), 4µL 1:25 diluted primary PCR product, 0.2µL Platinum Taq Polymerase, and 37.3µL H_2_O for a total volume of 50µL. Cycling conditions were the same as the primary PCR but 20 cycles for step 2. Samples were then pooled and submitted to the University of Iowa Institute of Human Genomics for sequencing on the Illumina NovaSeq 6000.

MAGeCK was used to identify kinases that show significant depletion with trametinib treatment (p ≤ 0.05, log fold-change ≤ -1).

### Western blotting

Protein lysates were harvested using 1X RIPA (Thermofisher #89900) supplemented with cOmplete™, Mini Protease Inhibitor Cocktail (Sigma Aldrich #11836153001) following the Thermofisher RIPA Lysis and Extraction Buffer protocol. NuPage 4-12 % B-T 1.5 mm × 15 well, (Thermofisher NP0323BOX) gels were loaded with ∼20µg protein and run at 100V for 1.5 hours. Transfer was performed at 18V overnight, and membranes were blocked in AquaBlock Blocking buffer (Arlington Scientific #PP82P). Primary antibody incubation was performed at 4^°^C overnight. Primary antibodies were as follows: α-tubulin (ab6160), Erk1/2 (CST 9102), pErk1/2 (T202/Y204) (CST 9101), Akt (BD Bioscience 610861), pAkt (S473) (CST 4060), Fak (CST 71433), pFak (Y397) (Thermofisher 44-624G), Mek1/2 (CST 4694), pMek (S298) (ab96379). Secondary antibody incubation was performed for 1 hour at room temp. Secondary antibodies included: Goat anti-Mouse IgG (Invitrogen A21058), Goat anti-Rabbit IgG (Invitrogen A11369), anti-Rat (Biotium 20096-1).

### Flow cytometry

Cytopainter (#ab176736) was used for cell proliferation staining following the protocol: dilute Cytopainter to 1X in HHBS buffer (1X HBSS (Gibco #14025-092) supplemented with 20mM HEPES buffer (Gibco #15630-080)), incubate for 30 minutes at 37^°^C, wash with HHBS buffer, and proceed with experimental setup or flow cytometry.

Apoptotic cells were stained with FITC Annexin V (Biolegend #640906) following the protocol: wash with Biolegend cell staining buffer (#420201), resuspend in HEPES buffer at ∼1 million cells/mL, add FITC Annexin V (2.5µg/mL) to 100µL cells, incubate at room temp for 15 minutes (protected from light), add 400µL HEPES buffer, and then analyze by flow cytometry. Staurosporin (Millipore Sigma #S6942) was used as a positive control to induce apoptosis at 1µM for 24 hours. Flow was performed at the University of Iowa Flow Core.

## FIGURES AND TABLES

**Figure S1.**
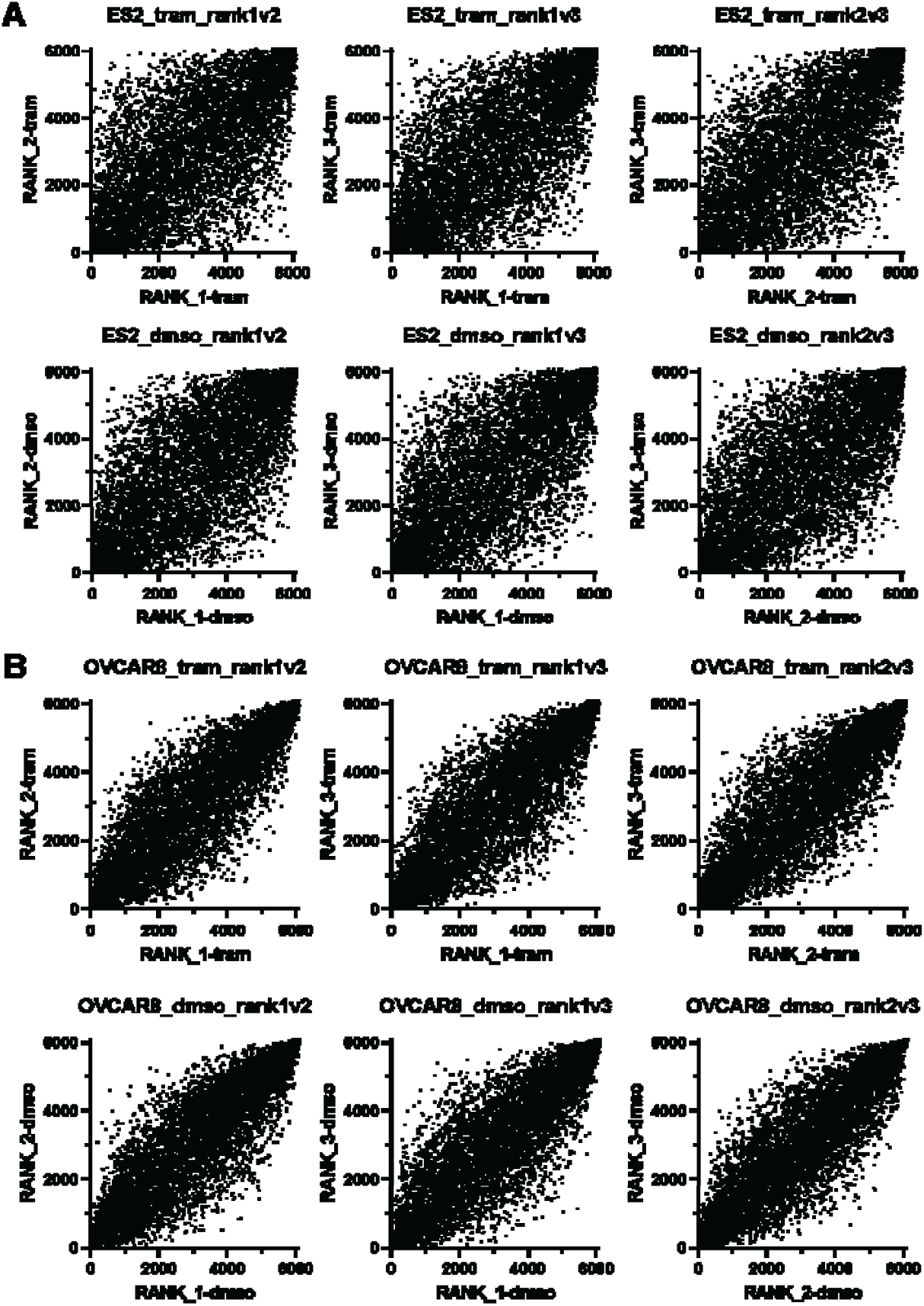
Spearman scores for targeted CRISPR kinome screen. (A) ES-2 sgRNAs ranked and compared between biological replicates (top, trametinib treated cells; bottom, vehicle treated cells). (B) OVCAR-8 sgRNAs ranked and compared between biological replicates (top, trametinib vehicle treated cells; bottom, treated cells).

**Figure S2.**
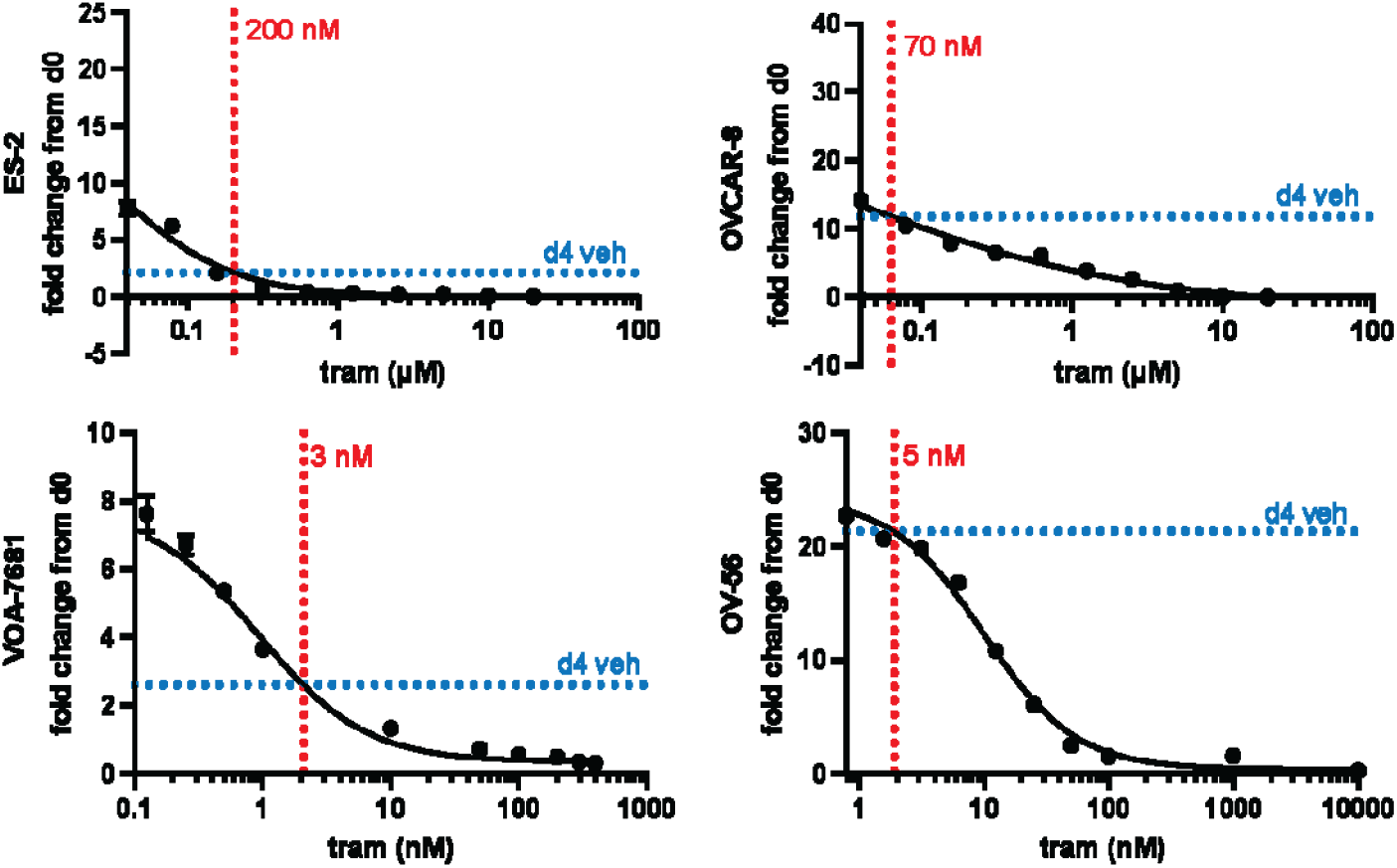
Dose determination for the CRISPR kinome screen based on 10-day dose response curves. Doses were determined based on the 10-day dose response curves generated for (A) ES-2, (B) OVCAR-8, (C) VOA-7681, and (D) OV-56. The horizontal blue dotted line indicates the fold change of vehicle-treated cells after 4 days. The vertical red dotted line indicates the dose of trametinib at which 10 days of treatment would yield a similar fold change as the 4 days of vehicle treatment. Vehicle and trametinib treated cell populations were collected after the same number of population doublings.

**Table S1: list of hits from the kinome screen**

[need to generate an excel file with only necessary information]

**Table S2.** Synergy scores for each tested combination therapy and cell lines.

| Cell line | drug1 | drug2 | ZIP<br>synergy | ZIP<br>p-value | HSA<br>synergy | HSA<br>p-value | Loewe<br>synergy | Loewe<br>p-value | Bliss<br>synergy | Bliss<br>p-value |
| --- | --- | --- | --- | --- | --- | --- | --- | --- | --- | --- |
| ES-2 | avutometinib | VS-4718 | 1.1764 | 2.46E-01 | 3.6631 | 3.63E-02 | 3.1876 | 3.22E-02 | -0.5864 | 6.93E-01 |
| ES-2 | trametinib | capivasertib | 23.2075 | 2.83E-10 | 19.4635 | 3.01E-08 | 18.9712 | 1.51E-07 | 24.4224 | 1.71E-10 |
| ES-2 | avutometinib | capivasertib | 9.3548 | 8.83E-11 | 10.4215 | 4.09E-05 | 6.9597 | 4.51E-04 | 8.9388 | 6.46E-08 |
| OVCAR-8 | avutometinib | VS-4718 | 10.5675 | 1.41E-09 | -2.3513 | 6.49E-02 | -4.5439 | 2.09E-03 | 13.5684 | 1.25E-07 |
| OVCAR-8 | trametinib | capivasertib | 4.9028 | 2.32E-06 | 7.9122 | 7.40E-08 | 6.4056 | 2.49E-07 | 4.8624 | 5.77E-05 |
| OVCAR-8 | avutometinib | capivasertib | 3.5113 | 1.56E-06 | -0.9167 | 3.61E-01 | -2.0134 | 4.94E-02 | 3.7591 | 5.32E-04 |
| OV-56 | avutometinib | VS-4718 | 2.9896 | 9.02E-04 | 8.4295 | 1.81E-07 | 6.2939 | 6.98E-06 | 2.2058 | 3.66E-02 |
| OV-56 | trametinib | capivasertib | 7.5938 | 1.35E-06 | 11.4604 | 4.33E-08 | 9.7368 | 1.04E-08 | 7.4605 | 2.68E-06 |
| OV-56 | avutometinib | capivasertib | 9.3548 | 8.83E-11 | 10.4215 | 4.09E-05 | 6.9597 | 4.51E-04 | 8.9388 | 6.46E-08 |
| VOA-7681 | avutometinib | VS-4718 | 2.4711 | 1.88E-02 | 3.8832 | 9.63E-03 | 1.8240 | 5.53E-02 | 2.0000 | 1.29E-01 |
| VOA-7681 | trametinib | capivasertib | 12.1596 | 1.24E-12 | 15.5933 | 2.48E-11 | 12.6120 | 4.29E-11 | 12.1461 | 3.64E-11 |
| VOA-7681 | avutometinib | capivasertib | 8.7631 | 2.48E-15 | 13.5711 | 5.38E-12 | 11.1519 | 6.60E-12 | 8.4229 | 1.02E-11 |

**Table S3.**
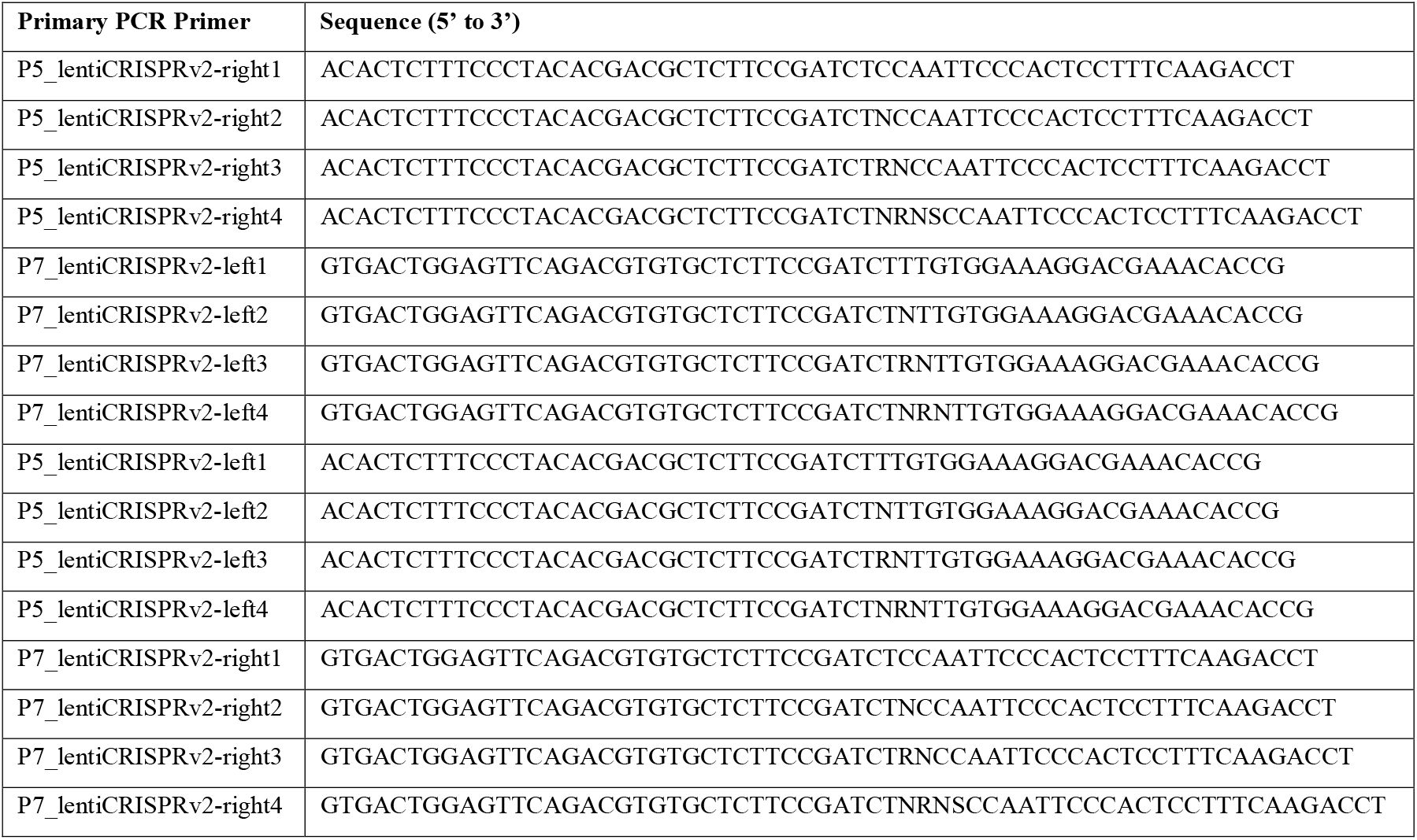
Primer sequences for CRISPR kinome screen library preparation.

